# Pairwise comparisons across species are problematic when analyzing functional genomic data

**DOI:** 10.1101/107177

**Authors:** Casey W. Dunn, Felipe Zapata, Catriona Munro, Stefan Siebert, Andreas Hejnol

**Affiliations:** Department of Ecology and Evolutionary Biology, Brown University, Providence, RI, USA; Current address: Department of Ecology and Evolutionary Biology, University of California Los Angeles, Los Angeles, CA, USA; Current address: Department of Molecular & Cellular Biology, University of California at Davis, Davis, CA, USA; Sars International Centre for Marine Molecular Biology, University of Bergen, Bergen, Norway

## Abstract

**Abstract:** There is considerable interest in comparing functional genomic data across species. One goal of such work is to provide an integrated understanding of genome and phenotype evolution. Most comparative functional genomic studies have relied on multiple pairwise comparisons between species, an approach that does not incorporate information about the evolutionary relationships among species. The statistical problems that arise from not considering these relationships can lead pairwise approaches to the wrong conclusions, and are a missed opportunity to learn about biology that can only be understood in an explicit phylogenetic context. Here we examine two recently published studies that compare gene expression across species with pairwise methods, and find reason to question the original conclusions of both. One study interpreted pairwise comparisons of gene expression as support for the ortholog conjecture, the hypothesis that orthologs tend to be more similar than paralogs. The other study interpreted pairwise comparisons of embryonic gene expression across distantly related animals as evidence for a distinct evolutionary process that gave rise to phyla. In each study, distinct patterns of pairwise similarity among species were originally interpreted as evidence of particular evolutionary processes, but instead we find they reflect species relationships. These reanalyses concretely demonstrate the inadequacy of pairwise comparisons for analyzing functional genomic data across species. It will be critical to adopt phylogenetic comparative methods in future functional genomic work. Fortunately, phylogenetic comparative biology is also a rapidly advancing field with many methods that can be directly applied to functional genomic data.

**Significance:** Comparisons of genome function between species are providing important insight into the evolutionary origins of diversity. Here we demonstrate that comparative functional genomics studies can come to the wrong conclusions if they do not take the relationships of species into account and instead rely on pairwise comparisons between species, as is common practice. We re-examined two previously published studies and found problems with pairwise comparisons that draw both their original conclusions into question. One study had found support for the ortholog conjecture and the other had concluded that the evolution of gene expression was different between animal phyla than within them. Our results demonstrate that to answer evolutionary questions about genome function, it is critical to consider evolutionary relationships.

## Introduction

The focus of genomic research has quickly shifted from describing genome sequences to functional genomics, the study of how genomes “work” using tools that measure functional attributes such as expression, chromatin state, and transcription initiation. Functional genomics, in turn, is now becoming more comparative– there is great interest in understanding how functional genomic variation across species gives rise to a diversity of development, morphology, physiology, and other phenotypes (1). These analyses are also critical to transferring functional insight across species, and will grow in importance in coming years.

Over the last three decades, a rich set of phylogenetic comparative methods has been developed to address the challenges and opportunities of trait comparisons across species (2–7). A central challenge is the dependence of observations across species due to the evolutionary history of species – more closely related species share many traits that evolved once in a common ancestor. This violates the fundamental assumption of observation independence in standard statistical methods. Phylogenetic comparative methods address this dependence. They have largely been applied to morphological and ecological traits, but are just as relevant to functional genomics (8). Even so, most comparative functional genomic studies have abstained from phylogenetic approaches and instead rely on multiple pairwise comparisons across species (Figure 1*A*). This leaves comparative functional genomic studies susceptible to statistical problems and is a missed opportunity to ask questions that are only accessible in an explicit phylogenetic context.

**Figure 1:**
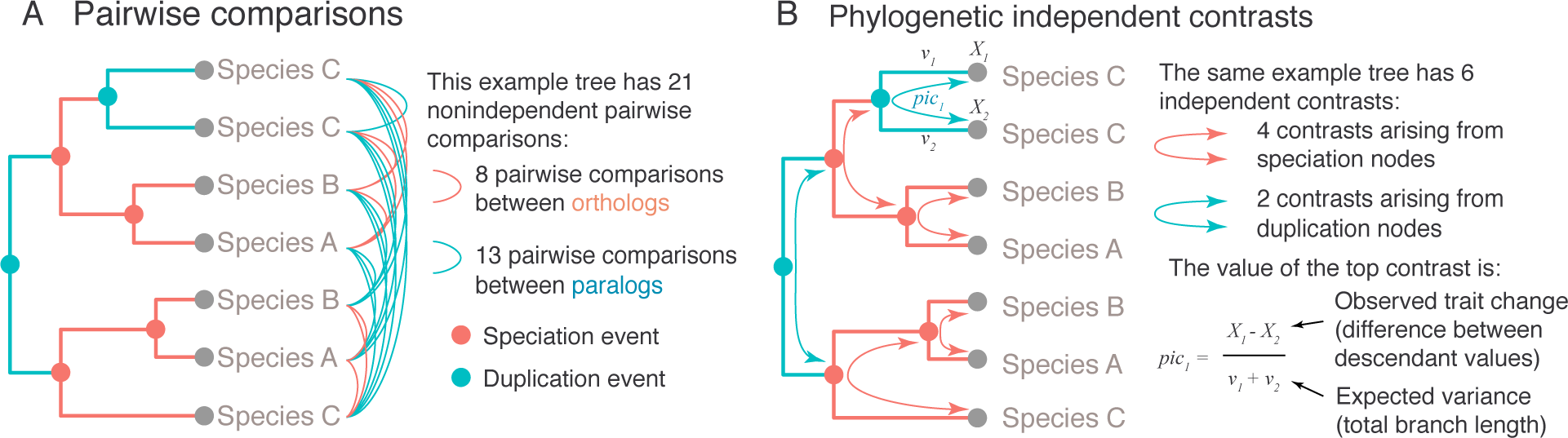
Pairwise and phylogenetic comparative approaches, illustrated on an example gene tree with multiple genes per species. The internal nodes of the tree are speciation and gene duplication events. (*A*) Many comparative functional genomic studies rely on pairwise comparisons, where traits of each gene are compared to traits of other genes across species. This leads to many more comparisons than unique observations, making each comparison dependent on others. (*B*) Comparative phylogenetic methods, including phylogenetic independent contrasts (2), make a smaller number of independent comparisons, where each contrast measures independent changes along different branches. Phylogenetic approaches are rarely used for functional genomic studies.

Phylogenetic comparative methods account for evolutionary history and explicitly model trait change along the branches of evolutionary trees (*e.g.*, Figure 1*B*). The value of these methods relative to to pairwise comparisons has been repeatedly shown in analyses of other types of character data (9–11). The application of these methods is illustrated by a study of leaf functional attributes for about 100 plant species (12). If phylogenetic relationships are not considered, analyses indicate a negative correlation of leaf lifespan with leaf size. With the application of phylogenetic comparative methods, however, this correlation disappears because most of the observed variance is due to differences between just two groups, conifers and flowering plants. Phylogenetic comparative methods capture this shift as change along a single branch deep in the tree, revealing that there is not a tendency of correlated change between these traits across the phylogeny. One reason that comparative functional genomic sudies have not embraced phylogenetic approaches is that there has not yet been a concrete demonstration that pairwise and phylogenetic comparative methods can lead to different results when considering functional genomic data. Here we examine this issue by re-evaluating the pairwise comparisons in two recent studies that compared gene expression across species.

The first study, Kryuchkova-Mostacci and Robinson-Rechavi (KMRR) (13), analyzed multiple vertebrate expression datasets to test the ortholog conjecture - the hypothesis that orthologs tend to have more conserved attributes (specificity of expression across organs in this case) than do paralogs (14). Using pairwise comparisons (Figure 1*A*), they found lower expression correlation between paralogs than between orthologs and interpreted this as strong support for the ortholog conjecture.

The second comparative functional genomic study we evaluate here is Levin *et al.* (16). This study analyzed gene expression through the course of embryonic development for ten animal species, each from a different phylum. Using pairwise comparisons, they found there is more evolutionary variance in gene expression at a mid phase of development than there is at early and late phases. They suggest that this supports an “inverse hourglass” model for the evolution of gene expression, in contrast with the “hourglass” model previously proposed for closely related species (17). Furthermore, they suggested that this provides biological justification for the concept of phyla. We previously described concerns with the interpretations of this result (18). Here we address the analyses themselves by examining the structure of the pairwise comparisons.

## Results and Discussion

### KMRR Reanalysis

#### Original pairwise test of the ortholog conjecture

KMRR (13) sought to test the ortholog conjecture. The ortholog conjecture (14) is the proposition that orthologs (genes that diverged from each other due to a speciation event) have more similar attributes than do paralogs (genes that diverged from each other due to a gene duplication event). The ortholog conjecture has important biological and technical implications. It shapes our understanding of the functional diversity of gene families. It is also used to relate findings from well-studied genes to related genes that have not been investigated in detail. It has be applied to many trait of genes, from gene sequence to biochemical properties to expression. While the ortholog conjecture describes a specific pattern of functional diversity across genes, it is also articulated as a hypothesis about the process of evolution– that there is greater evolutionary change in gene attributes following a duplication event than a speciation event.

Despite its importance, there have been relatively few tests of the ortholog conjecture. Previous work has shown that ontology annotations are not sufficient to test the ortholog conjecture (19, 20). Analyses of domain structure were consistent with the ortholog conjecture (21). There have been few tests of the ortholog conjecture with regards to gene expression (19), and KMRR is the most thorough such expression study to date.

KMRR considered several publicly available datasets of gene expression across tissues and species. Their expression summary statistic is Tau (22), an indicator of tissue specificity of gene expression. Tau can range from a value of 0, which indicates no specificity (*i.e.*, uniform expression across tissues), to a value of 1, which indicates high specificity (*i.e.*, expression in only one tissue). Tau is convenient in that it is a single number of defined range for each gene, though of course since the original expression is multidimensional this means much information is discarded. This includes information about which tissue expression is specific to. For example, if one gene has expression specific to the brain and another expression specific to the kidney, both would have a Tau of 1.

The KMRR analyses are based on pairwise comparisons (Figure 1*A*) between Tau within each gene family. Rather than make every pairwise comparison within each gene tree, they considered only a subset of pairwise comparisons in each particular analysis. They first selected a focal species, which varied from analysis to analysis. Ortholog comparisons were limited to pairs that include this species, and the only paralogs considered were those with the highest expression in this species. Note that this subset of pairwise comparisons still sample the same changes multiple times.

They found the correlation coefficient of Tau for orthologs to be significantly greater than the correlation coefficient of Tau for paralogs, *i.e.* orthologs tend to have more similar expression than do paralogs. From this they concluded that their analyses support the ortholog conjecture. They also concluded that this pattern provides support for a particular evolutionary process, that “tissue-specificity evolves very slowly in the absence of duplication, while immediately after duplication the new gene copy differs” (13).

#### Phylogenetic reanalyses

We reanalyzed the KMRR study using phylogenetic comparative methods. We focused on one of the datasets included in their analyses, that of Brawand *et al.* 2011 (24). This dataset is the best sampled in their analyses. It has gene expression data for six organs across ten species (nine mammals and one bird), eight of which were analyzed by KMRR and further considered here.

For each internal node in each gene tree, we calculated the phylogenetic independent contrast (2) (PIC) of Tau. This is the difference in values of Tau for descendant nodes scaled by the expected variance, which is largely determined by the lengths of the two branches that connect the node to its two descendants (Figure 1*B*). These contrasts were then annotated by whether each is made across a speciation or duplication event. The original description of independent contrasts (2) focused on assessing covariance between changes in two traits. Our use of contrasts is a bit different– we look for differences in evolutionary changes of one trait (differential expression) between two categories of nodes (speciation and duplication).

We mapped the Tau values calculated by KMRR (13) for the Brawand *et al.* 2011 (24) dataset onto 21124 gene trees parsed from ENSEMBL Compara (25). These are the same pre-computed trees that the orthology/ paralogy annotations KMRR used are based on. 8854 gene trees passed taxon sampling criteria (4 genes) after removing tips without Tau values and had at least one speciation event. Of these, 8513 were successfully time calibrated. These calibrated trees were used to calculate phylogenetic independent contrasts for 20945 duplication nodes and 67799 speciation nodes. One of these trees is presented in *SI Appendix* Figure 4 to demonstrate the analysis.

It is essential to have a null hypothesis that makes a distinct prediction from the prediction of the hypothesis under consideration. A suitable null hypothesis in this case is that there is no difference in the evolution of expression following speciation or duplication events (26). Under this hypothesis, we would predict that contrasts across speciation nodes and duplication nodes are drawn from the same distribution. Under the alternative hypothesis specified by the ortholog conjecture, that there is a higher rate of change following duplication events than speciation events, we would expect to see the distribution of duplication contrasts shifted to higher values relative to the speciation contrasts.

We did not find increased evolutionary change in expression following duplication events (Figure 2*B*). The Wilcoxon rank test does not reject the null hypothesis that the rate of evolution following duplications is the same as or less than the rate following speciation (p value = 1). Our phylogenetic comparative analysis, unlike the previously published pairwise comparative analysis (13), therefore finds no support for the ortholog conjecture in this system.

**Figure 2:**
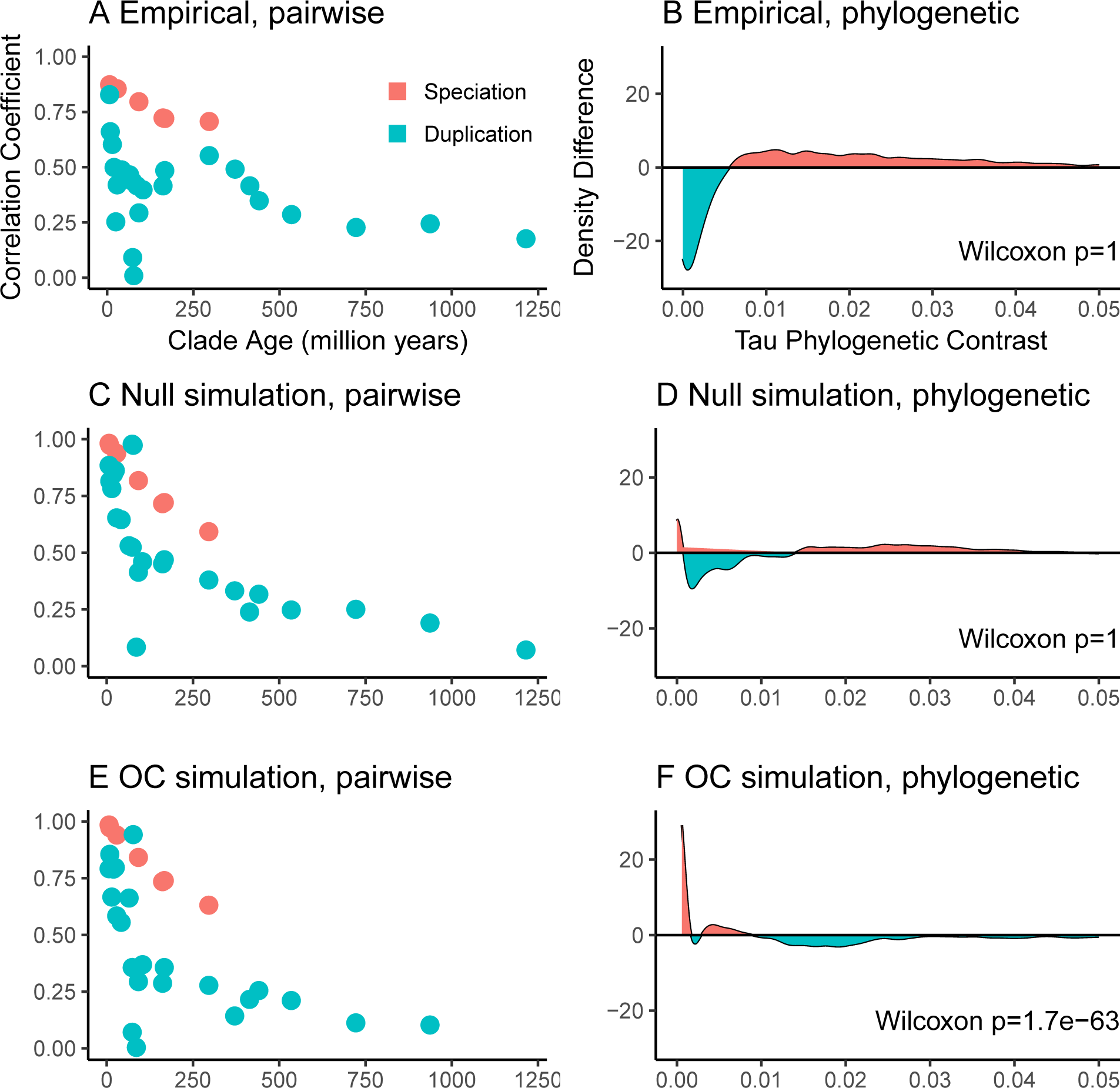
Pairwise (left column - *A, C, E*) and phylogenetic (right column - *B, D, F*) analyses of the original data (first row - *A, B*), data simulated under the null hypothesis (second row - *C, D*), and data simulated under the ortholog conjecture (third row - *E, F*). In the pairwise plots, each point indicates the correlation coefficient of Tau for a set of pairwise comparisons annotated with a specific node name (e.g, Primates) and event type (speciation or duplication, giving rise respectively to orthologs and paralogs). The phylogenetic plots show the difference between the density distributions for Tau phylogenetic contrasts for speciation and duplication events, where a value above 0 indicates an excess of speciation contrasts in the indicated interval. A horizontal line at 0 would indicate that the density distributions are identical. The top left pane (*A*) reproduces the pattern presented in Figure 2A of KMRR (13) of higher correlation across speciation events than duplication events, which they took as evidence of the ortholog conjecture. The recovery of a similar pattern under both simulations (*C, E*) indicates that it this pairwise approach does not make distinct testable predictions. The phylogenetic analysis of the original data (*B*) does not show an excess of larger contrasts for duplication events and does not reject the null hypothesis, providing no support for the ortholog conjecture. The bottom right panes validate the phylogenetic approach by showing that it does not reject the null when data are simulated under the null (*D*), but does reject the null when data are simulated under the ortholog conjecture (*F*).

We next examined the possibility that ascertainment biases were differentially impacting the inference of expression evolution following duplication and speciation events. Such a bias might obscure support for the ortholog conjecture. We focused on two possible sources of bias, node depth and branch length. We found no evidence that either affected our results (*SI Appendix*, Figure 5). We also examined the sensitivity of the results to the calibration times applied to speciation events on the gene trees. This is important because it is expected that genes from separate species have a common ancestor older than the time at which the species diverged from each other (27, 28). There is also uncertainty associated with the timing of these speciation events. We added random noise to the calibration times in replicate analyses, and all still failed to reject the null hypothesis (*SI Appendix*).

#### Understanding the incongruence between pairwise and phylogenetic methods

In order to better understand why our phylogenetic analysis supports a different conclusion (*i.e.*, no support for the ortholog conjecture) than the published analysis of KMRR (13) (*i.e.*, strong support for the ortholog conjecture), we first checked to make sure we could reproduce their result based on pairwise analyses. This is important since we are only looking at a subset of the data they considered, the Brawand *et al.* 2011 (24) dataset for gene trees that could be successfully time calibrated. In their Figure 1 (13), they present a higher Tau correlation coefficient between ortholog pairs than paralog pairs. We find the same here, with correlation coefficients of 0.75 for orthologs and 0.36 for paralogs.

Why is it that pairwise methods and phylogenetic methods lead to opposite conclusions? One reason is that multiple pairwise comparisons repeatedly sample the same evolutionary changes and in so doing violate statistical assumptions of independence, whereas phylogenetic comparative methods make multiple independent comparisons across non-overlapping branches of the tree. The other reason is that pairwise comparisons and phylogenetic comparative methods describe different things. Pairwise comparisons describe contemporary patterns, while phylogenetic methods infer historical processes (11). There need not be a different process of evolution following speciation and duplication for paralogs to be more different than orthologs. Any difference could be due to the structure of the gene phylogenies alone. If paralogs tend to be more distantly related to each other than orthologs, then there would be more time for differences to accumulate even if the rate of change is the same between the two. This is, in fact, the case for these data. While the mean distance (*i.e.*, total branch length) between orthologs is 333 million years, the mean distance between paralogs is 1473.8 million years. This is because the oldest speciation event is by definition the most recent common ancestor of the species included in the study, but many gene families underwent duplication before this time.

To test the hypothesis that ancient duplications that precede the oldest speciation event (*SI Appendix*, Figure 5*A*) impact the lower correlation of Tau between paralogs than between orthologs, we removed them. When we consider only the duplication events the same age or younger than the oldest speciation event (*SI Appendix*, Figure 5*B*), the paralog correlation coefficient increases from 0.36 to 0.56. This is much closer to the ortholog correlation of 0.75.

The KMRR study did investigate the impact of node age on correlation, but in a different way. In their Figure 2 (13), they grouped orthologs and paralogs according to the ENSEMBL node name of their most recent common ancestor, and plotted the correlation of Tau for each of these groups by the node age. They found that across the investigated range of node ages, ortholog pairs have higher Tau correlation than paralogs. We confirmed that we can replicate this result (Figure 2*A*). There are several difficulties with interpreting this plot, though. First, it does not just reflect the evolutionary processes that generated the data, it is also impacted by the phylogenies along which these processes acted. The expected covariance of traits that evolve under neutral processes is in fact defined by the phylogeny (3). Second, the correlation for each group is based on multiple non-independent pairwise comparisons.

To better understand this plot (Figure 2*A*), we performed simulations of Tau on the calibrated gene trees and then regenerated the figures. We did not modify the gene tree topologies or their inferred histories of duplication and loss. First, we simulated the evolution of Tau under the null model that it evolves at the same rate following duplication and speciation events. Under the null model, the plot of correlation coefficient to node age (Figure 2*C*) is very similar as for the observed data (Figure 2*A*). As in the original study, there is higher correlation coefficient across orthologs (0.74) than paralogs (0.31) when not considering node age. Phylogenetic analysis of the data simulated under the null hypothesis (Figure 2*D*) do not reject the null hypothesis (Wilcoxon p = 1), as expected.

We next simulated the evolution of Tau under the ortholog conjecture, where the rate of evolution of Tau following duplication was 2 fold the rate following speciation. The pairwise results of this heterogeneous model (Figure 2*E*) are nearly indistinguishable from the results under the null model (Figure 2*C*), and also have a higher correlation coefficient for orthologs (0.76) than paralogs (0.22). The phylogenetic analysis of the ortholog conjecture simulation (Figure 2*F*) does reject the null hypothesis (Wilcoxon p = 1.7 × 10^−63^).

These simulations have several implications. The pairwise comparisons used by KMRR cannot distinguish between the null hypothesis and ortholog conjecture. The pairwise results are strikingly similar under both hypotheses (Figure 2*C*, 2*E*). These simulations also serve to validate the phylogenetic methods applied to this problem. As expected, our phylogenetic analysis of independent contrasts does not reject the null hypothesis when data are simulated under the null model, and does reject the null hypothesis when the data are simulated under the ortholog conjecture. In contrast to pairwise methods, the phylogenetic analyses can test explicit predictions based on hypotheses about evolutionary process.

Since greater rates of expression evolution following duplication do not explain the lower correlation of Tau values in the pairwise comparison plots (Figure 2 *A, C, E*), this pattern must reflect some other property that is shared across these analyses. We found that the lower correlations for duplication comparisons can largely be explained by the greater variance in the age of duplication nodes. Each point in the pairwise correlation plot (Figure 2 *A, C, E*) summarizes multiple pairwise comparisons across nodes of a given event type (speciation or duplication, as indicated by color) and clade in the species tree (Theria, Mammalia, Amniota, etc…, as indicated by the age of the clade along the x axis). These clade names in turn are derived from the node annotations in the ENSEMBL Compara trees, which apply clade names to both speciation and duplication nodes. While speciation nodes in the gene trees have a clear correspondence to clades in the species trees, duplication nodes do not since duplication can occur at any point along any branch in the species tree. Neighboring clade ages are therefore a very rough approximation of the age of duplication events. This is apparent when node ages derived from calibrated trees are plotted against the age of the clade annotations for each node (*SI Appendix*, Figure 6*A*). Because duplication events that are all annotated with the same clade name can occur at very different times, pairwise comparisons across these nodes capture evolutionary changes in Tau across very different different branch lengths. They therefore have lower correlation than pairwise comparisons made across speciation nodes. Of the 956763 duplication nodes considered here, 263420 have a node age (as determined by the time calibrated trees) within 10% of their clade annotation. If only these duplication events closest to their annotated clade nodes are retained, the pattern of lower correlation of Tau evolution for duplication events disappears for the dataset simulated under the null model (*SI Appendix*, Figure 5*B*). This correction, though, comes at the high cost of discarding 72.5% of duplication nodes.

#### Implications for the ortholog conjecture

There has been considerable recent interest in, and controversy about, the ortholog conjecture (14, 15, 26, 29, 30). While some studies have presented support for the ortholog conjecture, our results are consistent with multiple studies that have not (14, 26, 31).

Our results suggest, at a minimum, that the ortholog conjecture is not a dominant pattern that is central to explaining the evolution of phenotypic diversity in gene families. This suggests that an alternative “neutral conjecture”, *i.e.* the conjecture that the evolution of gene traits tends to be the same following gene duplication and speciation events, may better explain the process and patterns of most gene evolution. Under this neutral conjecture, phylogenetic distance is a better predictor of the similarity of gene function than is the history of gene duplication and speciation. The ortholog conjecture does not have to be an all or nothing question, though. It may be the case that the rates of phenotypic evolution following duplication may be greater than that following duplication in some organisms, gene families, and evolutionary processes (29). We just do not find evidence for it when summarizing gene expression across tissues with Tau in these organisms. This calls into question the general predictive power of the ortholog conjecture with respect to gene expression, and until these processes are better understood it will be necessary to test for it in each situation. These tests should be articulated in terms of clear alternative hypotheses (26) that make distinct phylogenetic comparative predictions.

Lack of support for the ortholog conjecture has important biological implications. It indicates that the mechanism of gene divergence (speciation versus duplication) may not have as strong an impact on phenotypic divergence as sometimes proposed. It also has important technical implications. Having information on whether two genes are orthologs or paralogs provides little added information about expression beyond knowing how distantly related the two genes are. Rather than focus on whether genes are orthologs or paralogs when attempting to predict function, it may be more effective to simply focus on how closely related or distantly related they are. Closely related paralogs, for example, may tend to have more similar phenotypes than more distantly related orthologs (26).

This is also an example of the limitations of the concepts of orthology and paralogy (32). These terms can have straightforward meaning in small gene trees with simple duplication/speciation histories, but the utility of the terms breaks down on larger more complex gene trees. Orthology and paralogy are annotations on the tips of the phylogeny that are derived from the structure of the tree and history of duplication and speciation at internal tree nodes. In this sense, orthology and paralogy are statements about the internals of the tree that are distilled into statements at the tips of the tree. Much is lost in the process, though. For most questions it is much more direct to focus on the structure of the tree and the inferred processes within the tree, such as which internal nodes are duplication or speciation events and how much change occurs along the branches.

### Levin *et al.* reanalysis

#### Original pairwise analyses of developmental gene expression

Levin *et al.* (16) analyzed gene expression through the course of embryonic development for ten animal species, each from a different clade that has been designated as having the rank of phylum. They arrived at two major conclusions. First, animal development is characterized by a well-defined mid-developmental transition that marks the transition from an early phase of gene expression to a late stage of gene expression. Second, this transition helps explain the evolution of features observed among distantly related animals. Specifically, they concluded that animals from different phyla exhibit an “inverse hourglass” model for the evolution of gene expression, where there is more evolutionary variance in gene expression at a mid phase of development than there is at early and late phases. Closely related animals have previously been described as having an hourglass model of gene expression, where evolutionary variance in expression is greater early and late in development than at the midpoint of development (17, 33). Levin *et al.* conclude that this contrast between distantly and closely related animals provides biological justification for the concept of phyla and may provide a definition of phyla.

Levin *et al.* (16) arrived at this conclusion by making multiple pairwise comparisons of ortholog expression data sampled throughout the course of embryonic development. For each species pair, they identified the orthologs shared by these species. This list of shared genes was different from species pair to species pair. They characterized each of these orthologs in each species as having expression that peaks in early, mid, or late temporal phase of development. They then calculated a similarity score for each temporal phase for each species pair based on the fraction of genes that exhibited the same patterns in each species. The distributions of similarity scores are plotted in their Figure 4d (16), and their Kolmogorov–Smirnov (KS) tests indicated that the early distribution and late distribution were each significantly different from mid distribution (P < 10^−6^ and P < 10^−12^, respectively). This is the support they presented for the inverse hourglass model.

#### Reexamination of pairwise comparisons

We examined the matrix of pairwise comparisons used as the base for the KS tests and Figure 4d in *Levin et al.* (16), and thus as support for the “inverse hourglass” model. We found several problems resulting from the use of multiple pairwise comparisons. The first problems are specific to this particular implementation of pairwise comparisons. We found that every data point was included twice because both reciprocal pairwise comparisons (which have the same values) were retained. For example, there is both a nematode to arthropod comparison and an arthropod to nematode comparison. As a consequence, there are 90 entries for the 45 pairwise comparisons, and by doubling the data the significance of the result appears stronger than it actually is. After removing the duplicate values, the p values are far less significant, 0.002 for the early-mid comparison and on the order of 10^−6^ for early-late. In addition, the test they used (KS test) is not appropriate for the hypothesis they seek to evaluate. The KS test does not just evaluate whether one distribution is greater than the other, it also tests whether the shape of the distributions are the same. In addition, the samples in this dataset are matched (i.e., for each pairwise comparison there is a early, mid, and late expression value), which the KS test does not take into account. The Wilcoxon test is instead appropriate in this case. When applied to the de-duplicated data, the significance of this test is 0.02 for the early-mid comparison and on the order of 10^−7^ for early-late.

Once we addressed the issues above with the implementation of pairwise comparisons, we were able to explore more general issues that can be a problem when making multiple pairwise comparisons between species. We found that all five of the lowest values in the mid phase distribution (Figure 3*A*) are for pairwise comparisons that include the ctenophore (comb jelly). When the nine pairwise comparisons that include the ctenophore are removed, there is no significant difference between the early phase and mid phase distributions (p = 0.14 for the early-mid comparison and p < 10^−5^ for the late-mid comparison) and no support for the inverse hourglass (Figure 3b). This highlights a well understood property of pairwise comparisons across species (2, 12): evolutionary changes along a given branch, like those along the ctenophore branch, impact each of the multiple pairwise comparisons that includes that branch. The pairwise comparisons are therefore not independent - different pairwise comparisons are impacted by changes along some of the same branches (Figure 1*A*). This can give the impression of a general pattern across the tree that is instead specific to changes along one part of the tree. The number of comparisons impacted by each change depends on the structure of the phylogenetic tree, *i.e.* how the species are related to each other. Phylogenetic comparative methods were developed specifically to address this problem (2).

**Figure 3:**
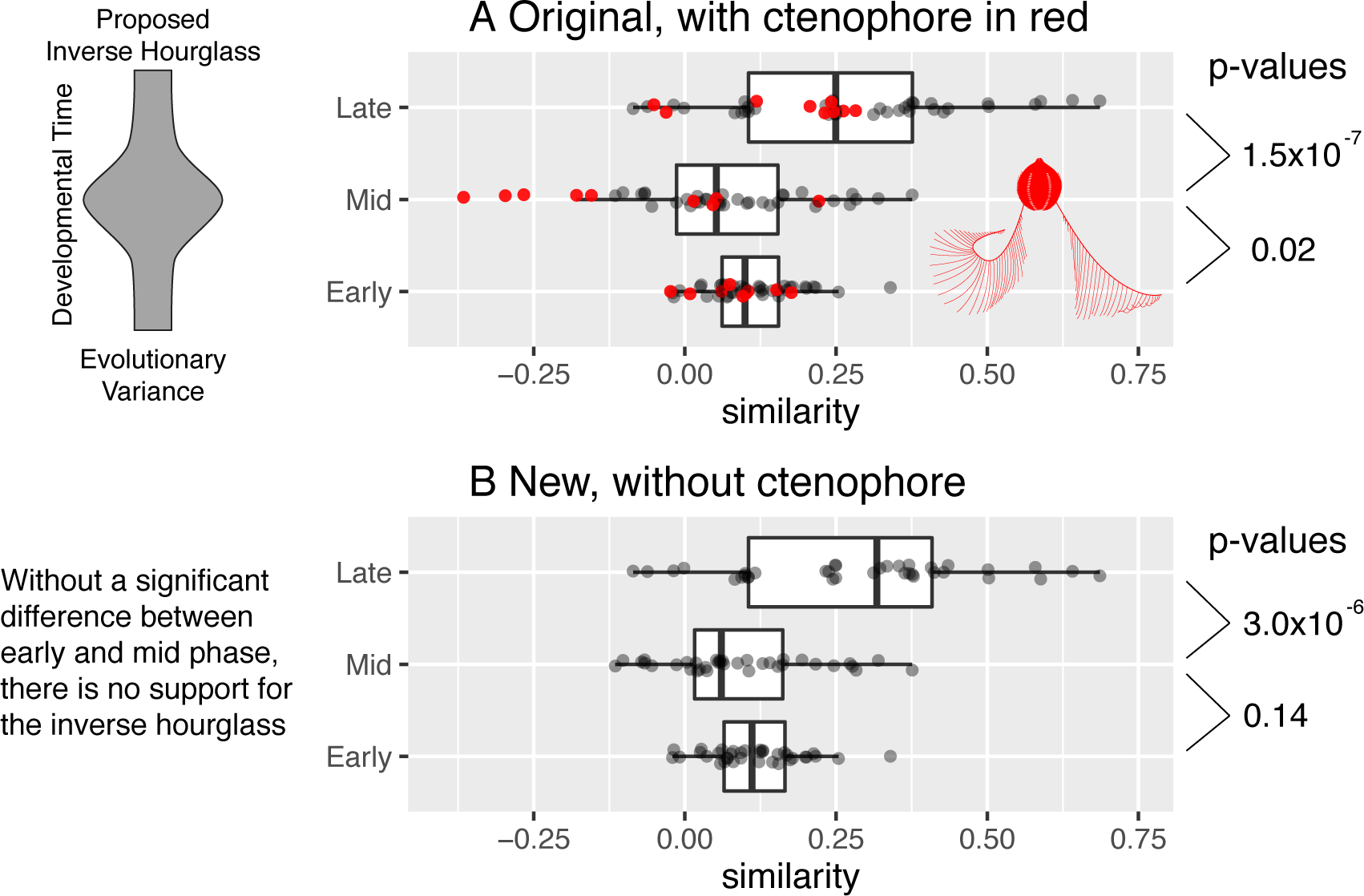
Distributions of pairwise similarity scores for each phase of development. Pairwise scores for the ctenophore are red. Wilcoxon test p-values for the significance of the differences between early-mid distributions and late-mid distributions are on the right. Model of variance, which is inversely related to similarity, is on the left. (*A*) The distributions as published by Levin *et al.* (16). Low similarity (*i.e.*, high variance) in the mid phase of development was interpreted as support for an inverse hourglass model for the evolution of gene expression. The five least-similar mid phase scores were all from the ctenophore. Published KS p-values, based on duplicated data, are in parentheses. The inset ctenophore image is by S. Haddock from phylopic.org. (*B*) The distributions after the exclusion of the ctenophore. The early and mid phase distributions are not statistically distinct.

While we demonstrate problems with pairwise comparisons that impacts the Levin *et al.* analysis, we did not perform a phylogenetic reanalysis of this study, as we did for the KMRR study. This is because the similarity metric computed in the pairwise comparisons of Levin *et al.* is not suitable for phylogenetic analysis. One issue with the similarity metric is that it is based on different genes for different species pairs, and therefore is not a trait whose evolution can be modeled across the phylogeny. A full phylogenetic reanalysis would be possible using upstream analysis products to re-derive new expression summary statistics.

#### Phylogenetic comparative methods in functional genomics

Our results highlight the importance of explicitly incorporating information about phylogenetic relationships when comparing functional genomic traits across species. Some of the most widely used phylogenetic comparative methods (2, 3) are already directly applicable to comparative functional genomic studies. There are also interesting new challenges at this interdisciplinary interface that will need to be addressed to fully realize the potential of phylogenetic comparative functional genomic studies. One such challenge is that most phylogenetic comparative analyses of covariance between traits have been developed to address problems with many more species (e.g., dozens or more) relative to the number of traits being examined. In comparative functional genomic analyses, there are often far fewer species because adding each species is still expensive, but high-throughput tools generate data for tens of thousands of traits per species. This creates statistical challenges as the resulting covariance matrices are singular and, if not treated appropriately, imply many false correlations that are artifacts of project design. We outlined these challenges and potential solutions in the context of gene expression elsewhere (34). In the same manuscript we also considered another issue of relevance here – the read counts generated by RNA-seq expression studies cannot be directly compared across species. This is because there are various species-specific technical factors that can be mistaken for differences in expression across species. These can be canceled out within species before making comparisons across species (34).

Recent advances in phylogenetic comparative methods are particularly well suited to addressing questions about the evolution of functional genomic traits. Most early phylogenetic comparative methods attempted to account for evolutionary signal to correct statistical tests for correlations between traits, while more recent methods tend to focus on testing hypotheses of evolutionary processes (35). The application of this newer focus to functional genomics provides an exciting opportunity to address long standing questions of broad interest, including the order of changes in functional genomic traits and shifts in rates of evolution of one functional genomic trait following changes in another trait.

We are not the first to apply phylogenetic comparative methods to functional genomic data. While the vast majority of comparative functional genomic studies have used standard pairwise similarity methods, a small number of comparative functional genomic studies have employed phylogenetic comparative approaches (36–38). For instance, a phylogenetic ANOVA (39) of the evolution of gene expression improves statistical power and drastically reduces the rate of false positives relative to pairwise approaches.

Addressing the statistical dependence of pairwise comparisons is not the only advantage of using phylogenetic comparative methods for functional genomic analyses. Another problem with the pairwise comparisons is that, except at the tips, they summarize changes along many branches in the phylogeny. Two paralogs that diverged from a duplication event deep in the tree may have many subsequent duplication and speciation events, and changes along all these branches will impact the final pairwise comparison. Phylogenetic methods have the advantage of isolating the changes under consideration (Figure 1*B*). Phylogenetic methods therefore avoid diluting the change that occurs along the branches that follow the node in question with changes along all subsequent branches. There may still be missing speciation events, due to extinction and incomplete taxon sampling, and missing duplication events, due to gene loss, but these omissions affect both methods.

## Conclusions

The fact that the first two comparative functional genomic studies we reanalyzed show serious problems with pairwise comparisons indicates that there are likely to be similar problems in other studies that use these methods. Future studies that compare functional genomic data across species will be compromised if they continue to use pairwise methods. Studies of evolutionary functional genomics should not be focused on the tips of the tree using pairwise comparisons. They should explicitly delve into the tree with phylogenetic comparative methods.

These analyses illustrate how important it is to not conflate evolutionary patterns with the processes that generated them. Finding a pattern wherein paralogs tend to be more different than orthologs is not evidence that there are different processes by which orthologs and paralogs evolve. This is also the expected pattern when they evolve under the same process but paralogs tend to be more distantly related to each other than orthologs are. The fact that multiple pairwise comparisons of developmental gene expression across diverse species share a particular pattern is not evidence of a general process that explains the differences between all species in the analysis. It is also the expected pattern when a single species has unique differences, and the evolutionary changes responsible for these differences are sampled multiple times in pairwise comparisons that span the same phylogenetic branches along which these differences arose. To use patterns across living species to test hypotheses about evolutionary processes it is also necessary to incorporate information about evolutionary relationships, *i.e.* phylogenies. There have been decades of work on building comparative phylogenetic methods that do exactly that, and they are just as relevant to comparing functional genomic traits across species as they are to comparing morphology or any of the other traits they are already routinely applied to.

## Methods

All files needed to re-execute the analyses presented in this document are available at https://github.com/caseywdunn/comparative_expression_2017. The most recent commit at the time of the analysis presented here was ed4802e4394e1fea838d934f52d873bc9e77eabe.

### KMRR reanalysis

The KMRR study (13) followed excellent practices in reproducibility. They posted all data and code needed to re-execute their analyses at figshare: https://figshare.com/articles/Tissue-specificity_of_gene_expression_diverges_slowly_between_orthologs_and_rapidly_between_paralogs/3493010/2. We slightly altered their Rscript.R to simplify file paths and specify one missing variable. This modified script and their data files are available in the github repository for this paper, as are the intermediate files that were generated by their analysis script that we used in our own analyses. We obtained the `Compara.75.protein.nh.emf` gene trees (25) from ftp://ftp.ensembl.org/pub/release-75/emf/ensembl-compara/homologies/ and include them in our github repository. These gene trees include branch lengths, annotate each internal node as being a duplication or speciation event, and provide a clade label for each internal node.

We considered only the data from Brawand *et al.* 2011 (24) for the eight taxa included in KMRR. We left in sex chromosome genes and testes expression data, which KMRR removed in some of their sensitivity analyses. This corresponded to the KMRR analyses that provided the strongest support for the ortholog conjecture and therefore the most conservative reconsideration of it.

After parsing the trees from the Compara file with treeio, which was recently split from ggtree (40), we added Tau estimates generated by the KMRR `Rscript.R` to the tree data objects. We then pruned away tips without expression data, retaining only the trees with 4 or more tips. We also only retained trees with one or more speciation events, as speciation events are required for calibration steps. This removes trees that have multiple genes from only one species after pruning away tips without expression data.

The gene trees were then time calibrated. The goal is not necessarily to have precise dates for each node, but to scale branch lengths so that they are equivalent across gene trees. This in turn scales the phylogenetic independent contrasts (which take branch length into account) so they can be compared appropriately. Before calibrating the trees, we had to slightly modify some of them. The node names in the ENSEMBL Compara (25) gene phylogenies are parsed from the NCBI Taxonomy database, which has many polytomies, rather than a bifurcating species phylogeny. One implication of this is that node names can be resolved in such a way that a speciation node can have the same name as one of its speciation node ancestors, as others have noted (41). If left unaddressed, this would force all intervening branches to have length zero and interfere with calibration. In particular, Hominini is the name for the clade that includes humans and chimps, while Homininae is the clade that includes humans, chimps, and gorillas. Because of the structure of the NCBI Taxonomy, both clades are labeled as Homininae in the Compara trees. To remedy this, we identified all clades labeled Homininae that have no gorilla sequence and renamed them Hominini. We then calibrated the trees by fixing the speciation nodes to the dates specified in the KMRR code, with the exception of Hominini and Homininae. These we set to 7 million years and 9 million years, drawing on the same TimeTree source (42) that KMRR used. We used the `chronos()` function from the R package ape (43) for this calibration, with the correlated model. See the *SI Appendix* for additional sensitivity analyses to time calibration. Some trees could not be calibrated with these hard node constraints, and were discarded.

For each node in the remaining calibrated trees, we calculated the phylogenetic independent contrast for Tau across its daughter branches with the pic() function in ape (43). We then collected the contrasts from all trees into a single table, along with other annotations including whether the node is a speciation or duplication event. This table, `nodes_contrast`, was then analyzed as described in the main text for the presented plots and tests.

### Levin et al. reanalysis

Levin *et al.* helpfully provided data and clarification on methods. We obtained the matrix of pairwise scores that underlies their Figure 4d and confirmed we could reproduce their published results. We then removed duplicate rows, applied the Wilcoxon test in place of the Kolmogorov-Smirnov test, and identified ctenophores as overrepresented among the low outliers in the mid-developmental transition column. An annotated explanation of these analyses is included in the git repository at https://github.com/caseywdunn/comparative_expression_2017/blob/master/levin_etal/reanalyses.md.

**Figure 4:**
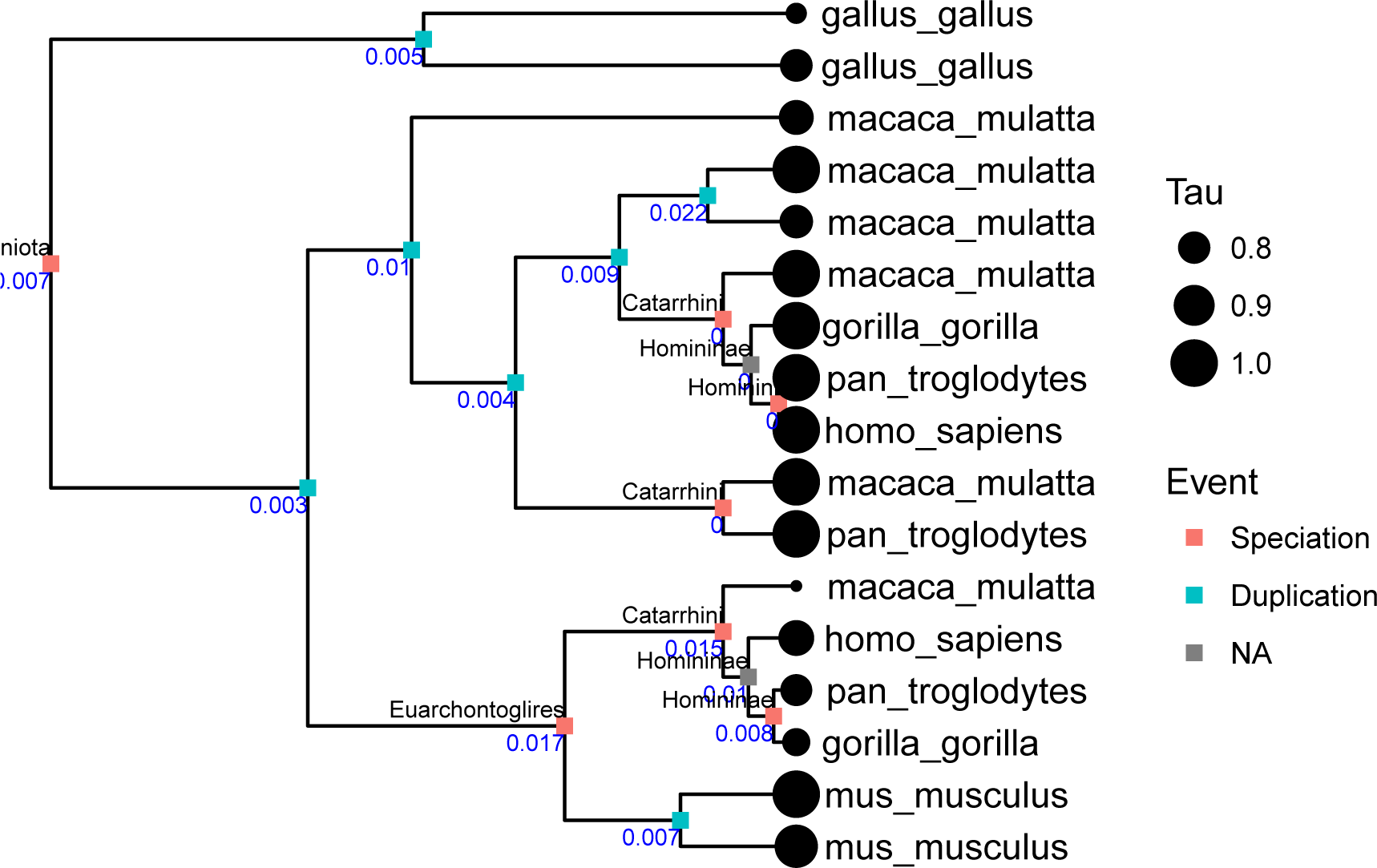
One of the gene trees from the phylogenetic analysis. The value of Tau (expression specificity) is indicated by the sizes of the circles at the tips of the tree. Whether an internal node is a speciation or duplication is indicated by color. Speciation nodes are labeled by clade name. Branch lengths are scaled to time. The blue number is the independent contrast for each node.

## Acknowledgments

Thanks to Steve Haddock, Alex Damian Serrano, August Guang, Bruno Vellutini, Zack Lewis, and Matthew Hahn for feedback on the manuscript. Marc Robinson-Rechavi provided very helpful information about the KMRR analyses and valuable suggestions for our manuscript. The Ensembl Comparative Genomics team, including Matthieu Muffato, provided excellent support for helping us better understand and use their Compara trees. CWD’s contribution was supported by the National Science Foundation (DEB-1256695 and the Waterman Award). AH was supported by the European Research Council Community’s Framework Program Horizon 2020 (2014–2020) ERC grant agreement 648861.

## Supplementary Information

### MRR Analyses

#### Summary statistics

**Table 1:**
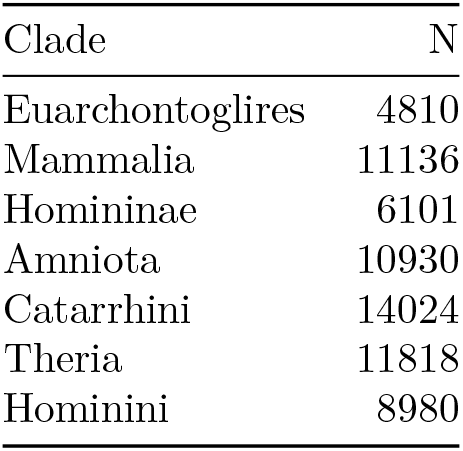
Number of speciation nodes N with contrasts by clade name.

| Clade | N |
| --- | --- |
| Euarchontoglires | 4810 |
| Mammalia | 11136 |
| Homininae | 6101 |
| Amniota | 10930 |
| Catarrhini | 14024 |
| Theria | 11818 |
| Hominini | 8980 |

**Table 2:**
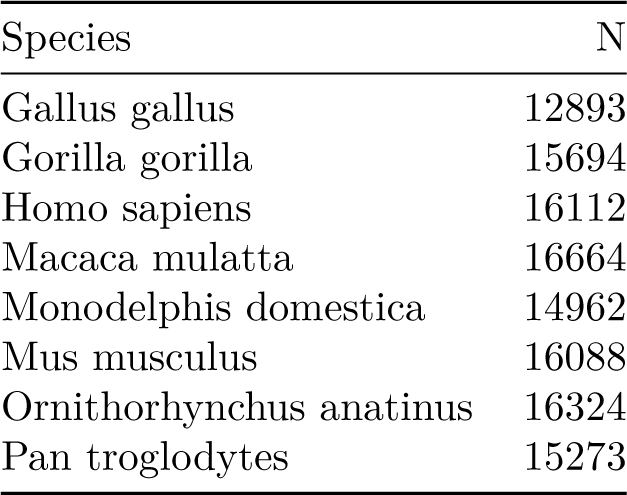
Number of tips N for each species in trees that were used to calculate the independent contrasts.

| Species | N |
| --- | --- |
| Gallus gallus | 12893 |
| Gorilla gorilla | 15694 |
| Homo sapiens | 16112 |
| Macaca mulatta | 16664 |
| Monodelphis domestica | 14962 |
| Mus musculus | 16088 |
| Ornithorhynchus anatinus | 16324 |
| Pan troglodytes | 15273 |

#### Investigation of potential ascertainment bias

While the age of speciation nodes is constrained, duplication nodes can be much older and therefore have a wider range of ages (*SI Appendix*, Figure 5*A*). This is because many gene duplication events are older than the most recent common ancestor of the species in the study. There are also technical factors that can lead to an excess of duplication events deeper in the tree. Gene tree estimation errors, for example, tend to lead to the overestimation of deep duplications (44). If independent contrast values also tended to to be lower at greater node depth, it could interact with the preponderance of duplications at greater depth to create a pattern of lower contrasts associated with duplication events. To test for such an effect, we remove duplication nodes that are older than the oldest speciation node. The general results are unchanged and this reduced dataset does not reject the null hypothesis that the rate of evolution following duplications is the same as or less than the rate following speciation (*SI Appendix*, Figure 5*B*).

**Figure 5:**
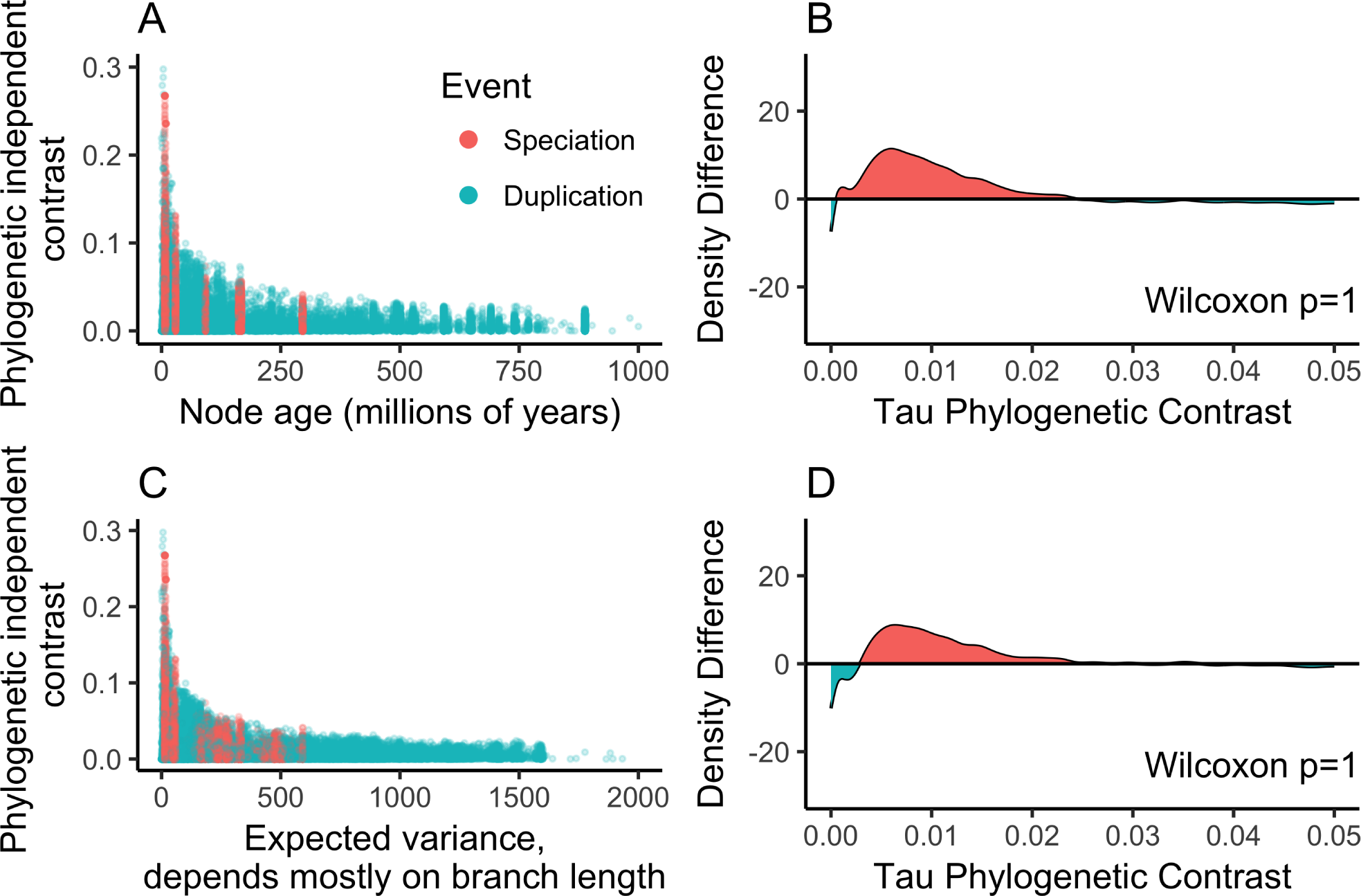
Investigation of possible ascertainment biases. (*A*) Magnitude of independent contrasts plotted against node age. Speciation nodes are calibrated to particular times, whereas duplication nodes have a wider range. (*B*) Difference between density distributions of contrasts for only the nodes that have an age less than or equal to the maximum age of speciation nodes. (*C*) Magnitude of independent contrasts plotted against expected variance, which is largely determined by branch lengths. Contrasts for speciation nodes have a narrower range of expected variance than do contrasts for duplication nodes. (*D*) Difference between density distributions of contrasts for only the nodes that have expected variance within the range of contrasts across speciation nodes.

**Figure 6:**
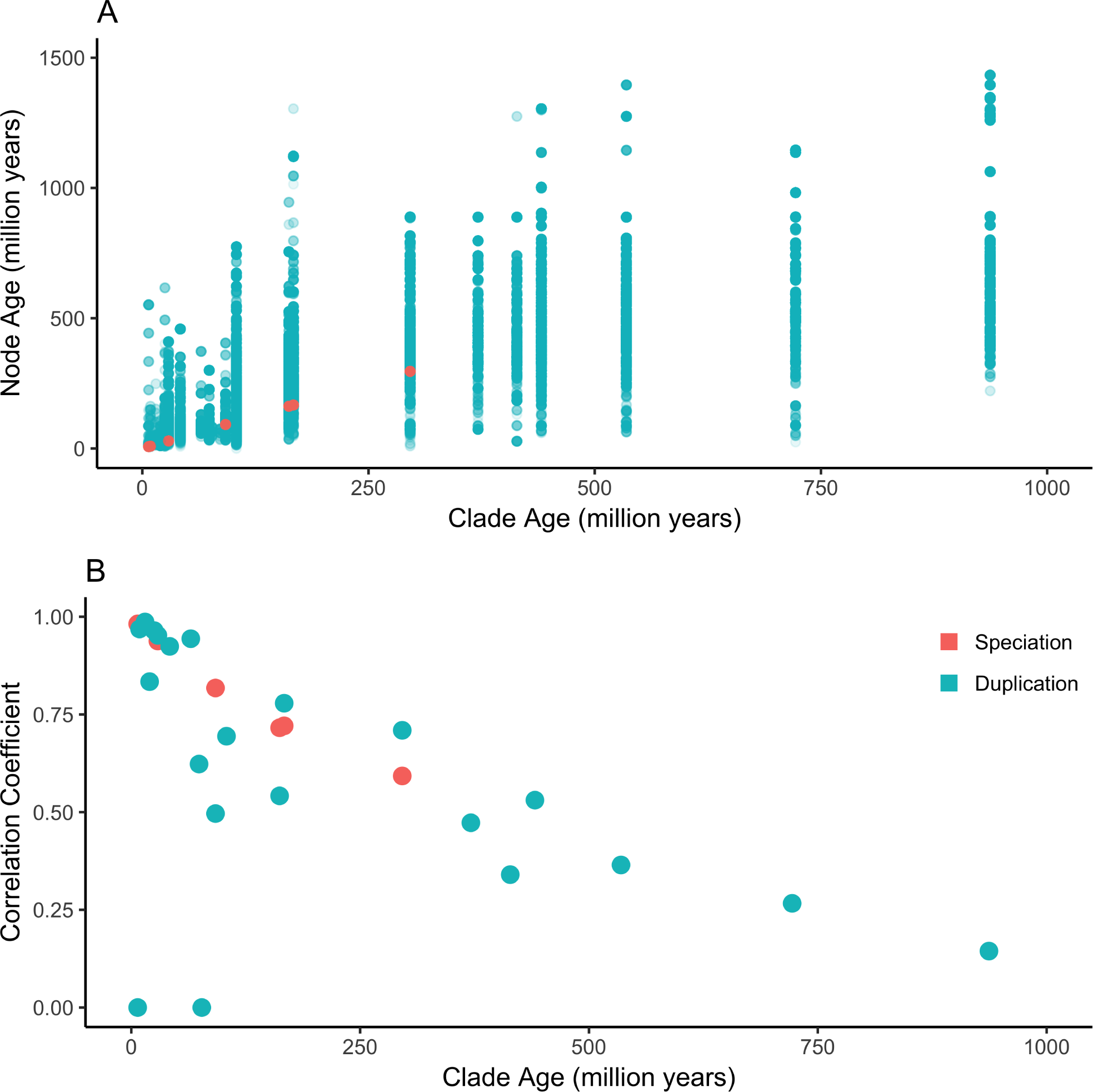
Origin of pairwise correlation plot structure. (*A*) Each speciation and duplication node in the COMPARA trees is annotated with a clade name, and the age of that clade (shown on the x axis) is used as the age of the pairwise comparisons (Figure 2 *A, C, E*, KMRR (13) Figure 2). The clade ages have a clear correspondence to the speciation node ages (red). Duplication (blue) ages for nodes assigned to the same clade, however, have large variation in calibrated node age (y axis). This means that summaries of expression differences across duplication events represent evolutionary changes that occurred over a wide range of time scales. (*B*) When duplication nodes whose calibrated node age deviates more than 10% from their annotated clade age are removed from the data simulated under the null distribution, duplication comparisons no longer have a uniformly lower expression correlation than speciation comparisons.

The independent contrast across a node is the amount of change observed between the daughter notes, scaled by the expected variance (2). The expected variance is principally determined by the lengths of the branches leading from the node to these daughters. The shorter the total length of the two branches leading to daughter nodes, the larger the contrast for a given observed difference. This is because the same difference across a shorter total branch length indicates a greater rate of evolutionary change. The expected variance of contrasts for speciation nodes is constrained by the branch lengths on the species tree, but the expected variance of contrasts for duplications has a much wider range (*SI Appendix*, Figure 5*C*). This could lead to biases if the lengths of branches that descend from duplication nodes tend to be overestimated. We therefore examined only the contrasts that fell within in the range of expected variance seen for speciation contrasts, excluding duplication contrasts that fall outside of this range. This reanalysis does not reject the null hypothesis either (*SI Appendix*, Figure 5*D*), indicating that branch length bias is not responsible for the result.

#### Investigation of sensitivity to calibration times

We examined the sensitivity of our results to the specification of calibration dates for the speciation nodes. In 10 reanalyses, we drew a new date for each calibration from a normal distribution with the mean of the original date and a standard deviation 0.2 times the original date. If any daughter nodes became older than their parent, we repeated the sampling until the dates were congruent with the topology. The minimum Wilcoxon p in these reanalyses was 1, *i.e.* none of them reject the null hypothesis that the rate of evolution of Tau is greater following duplication events than speciation events. This is consistent with the analysis that uses the calibration dates as specified, indicating that our results are robust to the selection of calibration times for speciation nodes.

#### Origin of pairwise correlation plot structure

#### Relationship between Tau and maximum expression

KMRR (13) Figure 3 differs from their other analyses in that it presents independent changes in expression between triplets of genes. Each triplet has two paralogs that arose from a duplication event and one unduplicated homolog. They find that there is a tendency for the paralog with the lowest expression to have the highest Tau. From this they conclude that following duplication there is a common trajectory of evolutionary change, where one paralog evolves to have lower expression and also become more tissue specific. There is, however, a negative relationship between Tau and maximum expression across all genes (*SI Appendix* Figure 7, *r*^2^ = 0.14, *slope* = 0.0519, *p* = 0). A simpler explanation for their plot is that it reflects the global tendency for Tau to decrease with maximum expression. The null expectation for any two genes sampled at random is for one to have higher maximum expression and lower Tau, and the other to have lower maximum expression and higher Tau. This global pattern could be due to both biological factors and the technical details of how Tau is calculated.

**Figure 7:**
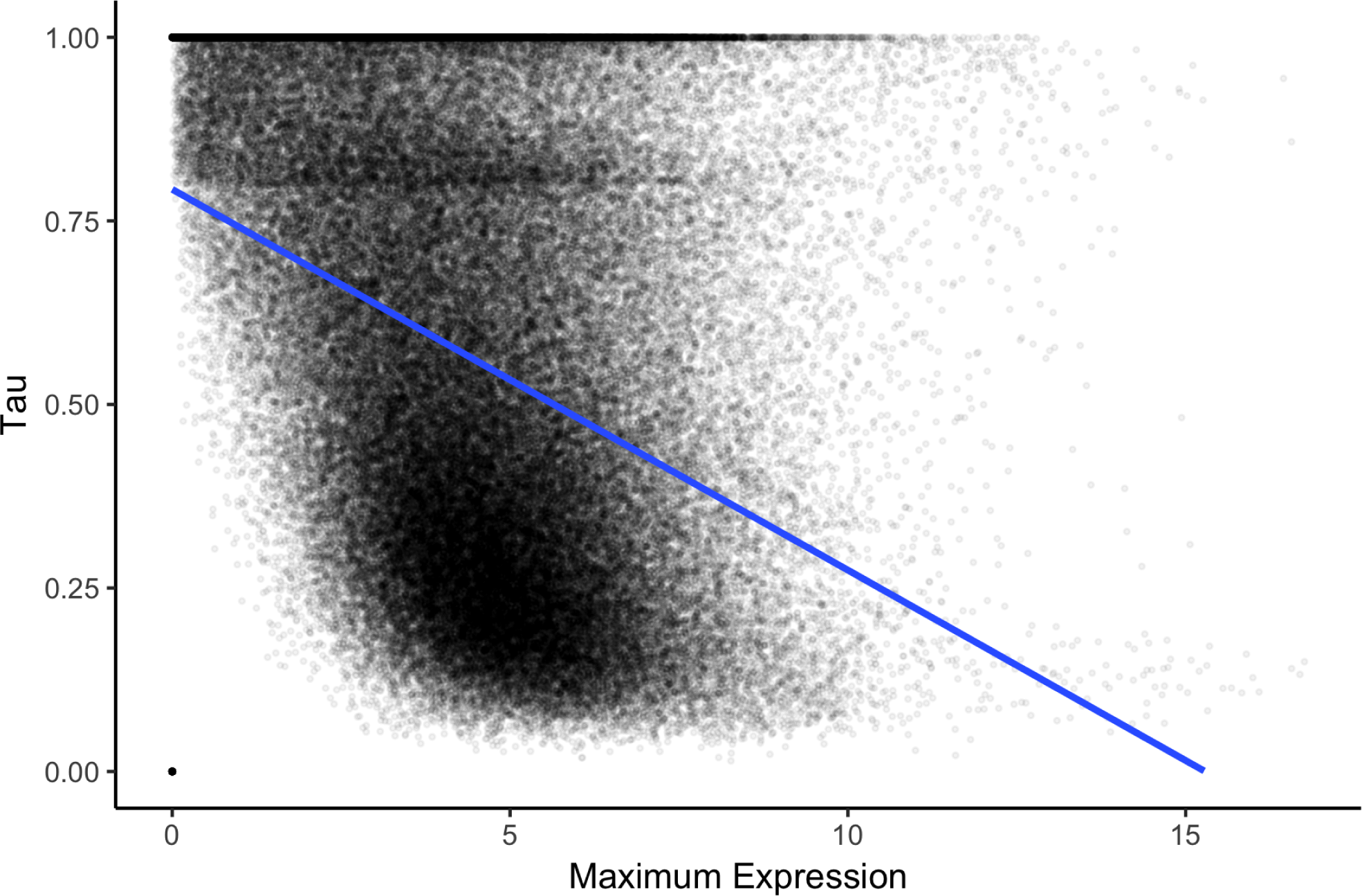
Genes with higher maximum observed expression tend to have lower Tau. The blue line is the linear model for the relationship between the two.

#### Software versions

This manuscript was computed on Thu May 04 20:38:30 2017 with the following R package versions.

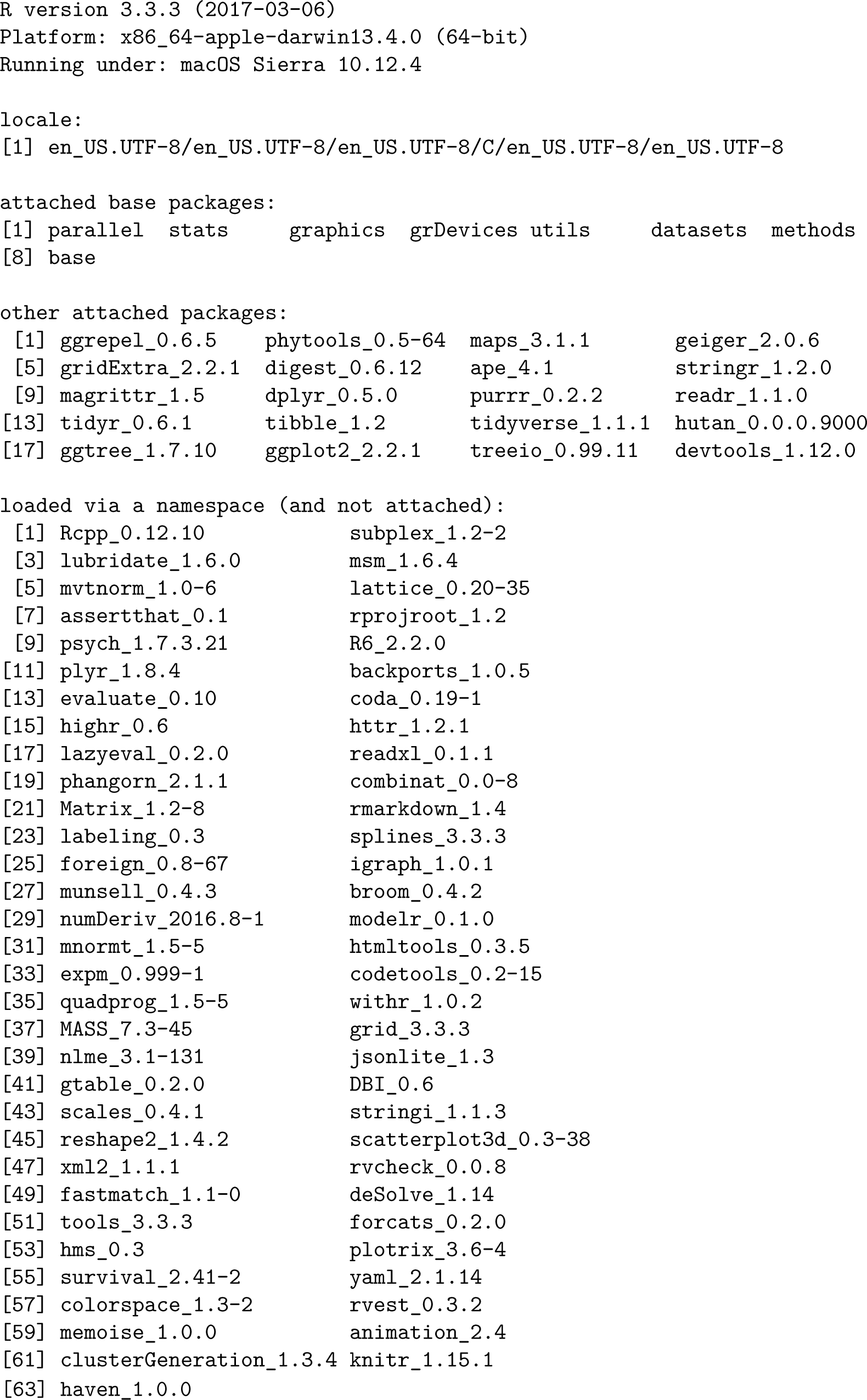

